# RNAse-free manufacture of Venezuelan Equine Encephalitis Virus (VEEV) plasmid DNA vaccine

**DOI:** 10.1101/2025.05.22.655656

**Authors:** Tyree Wilson, Michaela Harding, Charles Packninathan, Farhat Khan, Stasya Zarling, Sheetij Dutta

## Abstract

The long shelf-life and stability of DNA makes this platform highly attractive for low-cost, rapid delivery of pandemic response vaccines. Protocols utilized for clinical grade plasmid manufacture by contract development and manufacturing organizations are not readily accessible to academic and public research laboratories engaged in early-phase plasmid vaccine development. We present here the framework for DNA manufacturing using 3L-scale fermentation, anion-exchange chromatography and tangential flow filtration (TFF) leading to RNAase-free manufacture of plasmid DNA. The Venezuelan Equine Encephalitis Virus vaccine plasmid pWRG/VEEV, encoding the glycoprotein (E)3, E2, 6K and E1 genes was used as the prototype for this process development. The current effort yielded >95% pure and >80% supercoiled pWRG/VEEV plasmid preparations at 50-g wet cell weight scale. These data showed feasibility of manufacturing, highly pure pWRG/VEEV plasmid DNA using a cGMP compliant manufacturing process.

**Distribution Statement:** Approved for public release: distribution is unlimited

## Introduction

DNA delivery offers an unlimited potential for rapid scale-up and deployment of monovalent or polyvalent vaccines needed for pandemic response [1]. Human trials with DNA vaccines against viruses like Hantan, HIV, Flu, Zika, West Nile, Ebola and SARS-CoV-2 have shown a good safety profile [2-8]. Veterinary DNA vaccines against West Nile Virus, melanoma, Salmon pancreas disease and influenza are being investigated [9] and naked-DNA has been used for several gene therapy clinical trials [10, 11]. DNA has the advantage of thermostability over RNA, however immunogenicity of DNA encoded antigens remains low with only one DNA vaccine having received authorization for emergency-use against SARS-CoV-2 (ZyCov-D) [12]. DNA must cross the plasma membrane and the nuclear membrane for transcription and translation to occur, which requires a relatively high dose administration to elicit a productive immune response. Efforts to improve immunogenicity of DNA involve antigen modification, immune targeting sequences, co-delivery with lipid nanoparticles and the use of traditional vaccine adjuvants [13-15]. Economic strategies of manufacturing plasmid DNA are needed for rapid clinical evaluation of commercially viable DNA vaccines [13, 16-18].

Venezuelan Equine Encephalitis Virus (VEEV) is a mosquito-borne alphavirus that is prevalent in the Americas and causes periodic epidemics [19]. Although mortality is rare, infection with VEEV causes an encephalitic disease impacting the central nervous system. VEEV can also be aerosolized and therefore is a priority pathogen for the US Department of Defense [20]. While there is no FDA-approved vaccine for VEEV, the US Army developed the live-attenuated TC-83 and a formalin-inactivated C-84 vaccine to protect high-risk servicemen and lab entrants. TC-83 vaccine strain can be transmitted by mosquitoes and its adverse events profile and efficacy can be further improved. The C-84 vaccine mediates short-term protection and is used mainly to boost TC-83 non-responders. Efforts to develop safer vaccines led to exploration of DNA delivery of structural proteins from the VEEV 26S sub-genomic coding region (C-E3-E2-6K-E1) cloned into pWRG7077 DNA plasmid. The pWRG/VEE plasmid was highly immunogenic and protective in mice, guinea pigs and nonhuman primates [21].

The pWRG/VEE vaccine was originally produced under current cGMP environment by a contract manufacturer (Althea Technologies, Inc., San Diego, CA). Phase I trial (NCT01984983) at high or low dose pWRG/VEE showed that the product was safe and immunogenic [22, 23]. Another Phase I trial (NCT06002503) evaluating pWRG/VEE produced under cGMP by Aldevron, and delivered by needle-free jet injection, is ongoing. As improved versions of DNA vaccines emerge there will be need for rapid and cost-efficient production under cGMP environment. Plasmid DNA represents only 3% of bacterial cell mass and laboratory protocols for purification involve removing cellular RNA, genomic DNA and proteins using chemicals like cesium chloride, phenol and animal-derived enzymes (ribonuclease A), that are not recommended for large-scale manufacture of human products [24]. While there are several analytical reports on RNAse-free purification of plasmid DNA [25, 26], extensive modifications are required to adapt these methodologies to process development. In a feasibility study, fermentation, down-stream physical and anion exchange chromatography were optimized for biochemical and physical separation of plasmid from genomic DNA, proteins and RNA. The FDA recommended level of purity and supercoiled DNA content for vaccines [18, 27-32] was reached at 5 g- and 50 g- wet cell weight scale.

## Materials and methods

### Upstream production of cell mass

*E. coli* fermentations were initiated using a plant-based medium comprised of 1% Phytone, 0.5% Yeast Extract, 0.33% Ammonium Sulfate, 0.68% Potassium Phosphate monobasic, 0.024% Magnesium Sulfate, 0.71% Sodium Phosphate Dibasic, 0.5% glycerol. A starter culture (90 ml plant-based medium containing 100 mM potassium phosphate buffer (pH 7.2), 1% dextrose and 50 µg/mL kanamycin was inoculated with pWRG/VEE glycerol stock and grown at 210 rpm in 37°C for 12-16 hr in a shaking incubator. Fermentation was performed using a 3 L BioBlu 3f Macrosparge tank connected to an Eppendorf 320™ controller (Hamburg, Germany). The fermentation tank contained 2.6 L of culture medium, 300 mL 1 M Potassium Phosphate (pH 7.2), 75 mL 40% Dextrose, and 3 mL 50 µg/mL kanamycin. Fermentation conditions were set to 37°C, agitation 400 rpm, air-flow 3 SLPM, and pH 7.0. Dissolved oxygen was maintained at >30% saturation using agitation up to 800 rpm and oxygen cascade up to 0.5 SLPM. Cell growth (OD_600_) and glucose levels (GlucCell test strips; CESCO Bioengineering Co, Taichung, Taiwan) were monitored off-line, and as the dextrose level was depleted, a feed medium consisting of 80 g/L Dextrose and 40 g/L Yeast Extract was applied at 1 mL/min. A typical fermentation culture was harvested ∼30 hours post inoculation, in 1 L polypropylene bottles (Beckman Coulter, Indianapolis, IN, USA) by centrifugation at 6000 rpm for 20 min in 4°C. Cell pellet was stored at -20°C. Quality control during the fermentation process was performed by harvesting 1 ml culture for plasmid extraction using QIAprep spin columns (Qiagen, Germantown, MD, USA).

### Down-stream lysis and clarification

At 5g-scale, cells were resuspended in 25 mL **Resuspension buffer** (50 mM Tris, 10 mM EDTA; pH 8.0) by repeated pipetting. **Lysis Buffer-1** (41 ml; 0.96% NaOH) and **Lysis Buffer-2** (9 mL; 6% SDS) were added and cells were gently mixed by inversion and incubation at room temperature for 5 min. Lysate was neutralized by adding **Neutralization Buffer** (25 mL; 3 M Potassium Acetate), followed by gentle mixing by inversion and incubation on ice for 20 min. RNA was removed by adding an **RNA Reduction Buffer** (25 mL; 5 M Calcium Chloride) followed by gentle mixing by inversion and incubation for 20 min at room temperature. Cell debris were removed by centrifugation in 250 mL spin bottles at 8000xG for 10 min at 4°C, followed by sterile filtration of the supernatant using a 250 mL PES vacuum filter (Nalgene; 0.2-micron; Thermo Fisher Scientific, Waltham, MA, USA). At **50 g scale**, all reagent volumes shown above were proportionally increased 10-fold and clarification of the crude lysate was performed by a serial filtration using a Sartorius 50 µm PP3 Midicap filter, followed by Sartorius 1.2 µm PP3 Maxicap filter and Sartorius 0.8/0.2 µm 2 XLG filter (Sartorius, New Oxford, PA, USA). A peristaltic pump maintained at ∼60 mL/min was used for large-scale filtration. Cell lysate, once clarified, could be stored in 4°C overnight, prior to the diafiltration step.

### Diafiltration using Tangential Flow Filtration (TFF)

At 5g-scale, the lysate was passed through a 300 kDa MWCO Hollow Fiber Membrane (Midikross, Repligen Waltham, MA, USA; SA 115 Cm^2^, 41.5 cm; Part Number D02E30005S). At 50g-scale, TFF was performed using a 300 kDa Explorer 24-Reuse Hollow Fiber Membrane (1 mm x 320.6 cm^2^ long; 1/2” x 24”, Part Number WA-300, Göttengen, Germany). A Cole Palmer Peristaltic pump connected to Masterflex tubing (Thermo Fisher Scientific, Waltham, MA, USA) was assembled and an AISI pressure gauge was used to maintain column pressure (Millipore Sigma, Burlington, MA, USA). The TFF flow rate for 5g-scale runs was maintained at ∼40 mL/min and at ∼60 mL/min for the 50g-scale. Prior to TFF, the hollow fiber column was washed with 5 CV of water followed by 2 CV of **Equilibration buffer** (50 mM Tris, 400 mM NaCl; pH 8.0). The lysate was diluted in an equal volume of buffer and concentrated to final volume of ∼125 mL for 5g-scale and ∼1.2 L for 50g-scale. During TFF the pressure was maintained <10 PSI and 5 cycles of dilution and concentration were performed until a target pH (pH ∼8.0) and conductivity (∼40 mS/cm) were reached.

### Anion Exchange Chromatography

For the 5g-scale, 10 mL Fractogel DMAE was packed in a Vantage L Laboratory Column VL 11×250 cm column (Millipore Sigma, Burlington, MA, USA, Part Number 96100250), connected to an AKTA Pilot system (Cytiva Life Sciences, Marlborough, MA, USA). For 50g-scale, 35 ml Fractogel DMAE was packed in a Waters AP-2 column 20 mm x 100 mm (Waters Corporation, Milford, MA, USA). Columns were equilibrated with 5 CV **Equilibration Buffer** (4 mL/min for small scale or 20 ml/ml large scale, respectively), cleared lysate was loaded (1 ml/min or 10 ml/min) and flow-through was collected. The column was washed with 5 CV **Equilibration Buffer** (4 mL/min or 20 ml/min) followed by 5 CV **RNA Wash buffer** (50 mM Tris, 0.6 M NaCl; pH 8.0). The plasmid DNA was eluted in **Elution buffer** (50 mM Tris, 1 M NaCl; pH 8.0). OD_260_, pH, conductivity and column pressure were monitored.

### Isopropanol (IPA) precipitation

The peak elution fractions from anion exchange column were pooled and equal volume of **Isopropanol** was added, at room temperature, mixed by inversion, and stored -20°C, overnight. The plasmid was recovered by centrifugation at 14,000xg, 30 min at 4°C in Oak Ridge polycarbonate centrifuge tubes (Thermo Fisher Scientific, Waltham, MA, USA). The pDNA pellet were air dried for 60 min and resuspended in 1 ml (5g-scale) or 5 ml (50g-scale) **TE buffer** (10 mM Tris-HCl, 1 mM EDTA; pH 8.0).

### Qualitative analysis by agarose gel electrophoresis

DNA concentration was determined using Nanodrop 2000C (Thermo Fisher Scientific Waltham, MA, USA). Plasmid samples were analyzed by horizontal electrophoresis using 0.8% pre-cast agarose gels (E-Gel; Thermo Fisher Scientific Waltham, MA, USA) and scanned with a digital imager (Bio-Rad, Philadelphia, PA). Track-it DNA Ladder (Thermo Fisher Scientific Waltham, MA, USA) was loaded as molecular weight standard and cGMP grade pWRG/VEEV was used as the positive control.

### HPLC analysis

DNA purity was determined using CIMac pDNA-0.3 analytical column (Sartorius, New Oxford, PA, USA), as per manufacturer’s application note. The mobile phase consisted of **Solvent A** (0.2 M Tris; pH 8.0) and **Solvent B** (0.2 M Tris containing 1 M NaCl; pH 8.0), and the flow rate was maintained at 1.0 mL/min. Gradient conditions were as follows: 0-2 mins, 0% B; 2-3 min, 0-60% B; 3-6 min, 60% B; 6-16 min, 60-68% B; 16-17 min, 100% B; 17-20 min, 0% B. OD_260_ of the eluent was monitored. Prior to loading, samples were diluted to 100-150 ng/µL in Solvent A. pWRG/VEE produced under cGMP conditions and *E. coli* total RNA (1 mg/mL; Thermo Fisher Scientific, Waltham, MA, USA, PN: AM7940) were included as the controls.

### Residual protein analysis

Residual protein was estimated using NanoOrange^R^ Protein Quantitation kit (Molecular Probes, Eugene, OR, USA). Briefly, a standard curve was plotted between 10,000 to 300 ng/ml BSA. Sample was tested at 1:1000 and 1:25 dilutions according to the manufacturer’s protocol.

## Results

### Fed-batch fermentation (3L-scale)

Animal-free material was sourced to grow the pWRG/VEE bacterial clone in fermentation. The overnight culture grown in a shake-flask (final OD_600_ ∼4.0) was used to seed a 3L-scale fermentation (30-fold dilution; starting OD_600_ ∼0.1; starting glucose ∼12 gL^-1^). Cells grew linearly up to ∼9.0 OD within 5 hrs and to OD ∼20.0 at ∼20 hr post inoculation **(Figures 1A)**. A high glucose containing medium was fed to restore linear growth, reaching a final OD ∼40.0, at ∼30 hr post inoculation. The 3L fermentation yielded ∼45 g/L cell paste and the product identity an integrity were confirmed by agarose gel electrophoresis, sequencing **(Figure 1B; Figure S1)**.

**Figure 1.**
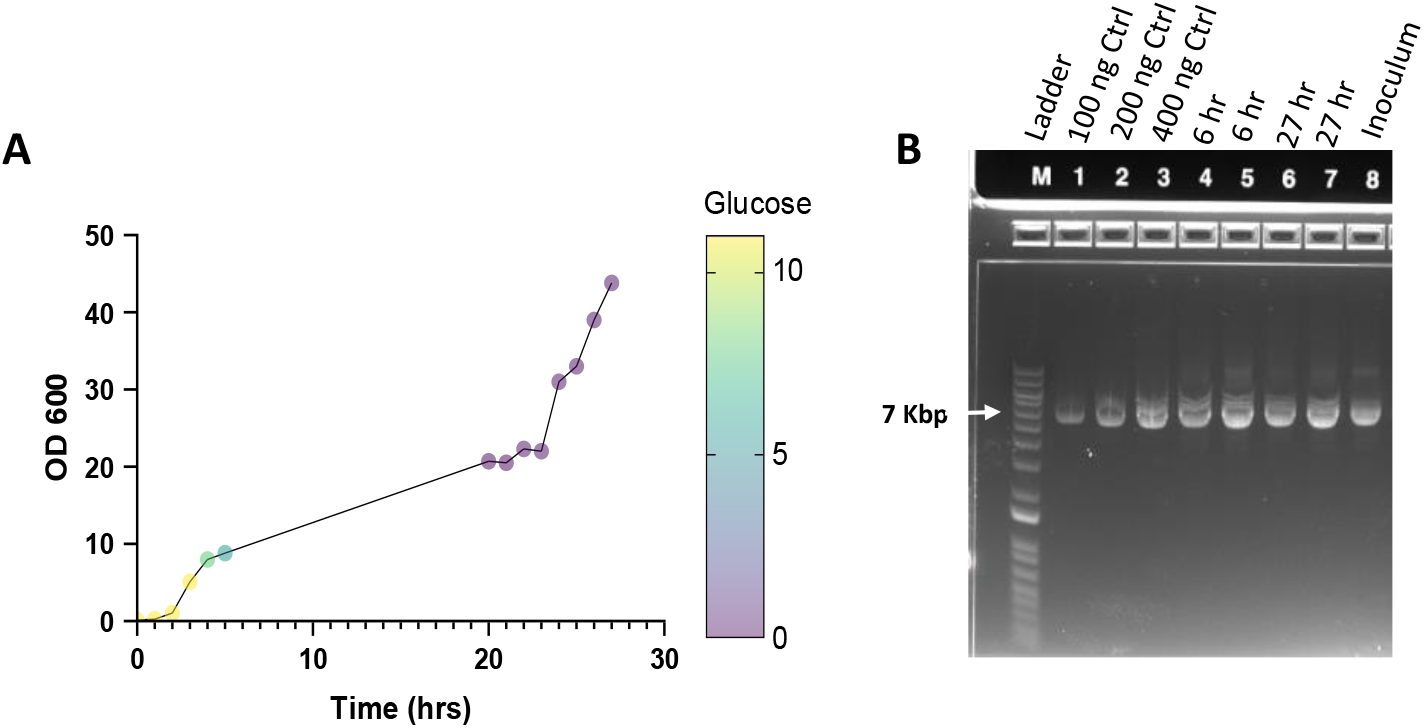
Fermentation at 3L-Scale: **(A)** Typical fermentation growth curve, OD600 nm *vs*. time (hrs) *vs*. glucose concentration (g/L). **(B)** Agarose gel analysis of the final product; Lane M, DNA ladder; lanes 1, 2, 3, control VEEV plasmid; lanes 4, 5, DNA from culture harvested at T=6 hr; lanes 6, 7, DNA from culture harvested at T=27 hr; lane 8, DNA from the shake-flask inoculum.

### Gravity column purification development

Small-scale purification runs using gravity columns were performed, at 5g-scale, to initially optimize the purification. Birnboim and Dolly alkaline lysis was followed by CaCl_2_ treatment [33] and tangential flow filtration (TFF), which retains molecules >4000 kDa. Small-scale purification experiments showed that CaCl_2_ addition was critical to reduce the RNA content **(Figure S2)** and four to five rounds of TFF dilution, concentration cycles, were optimal for conductivity and pH to be adjusted for plasmid DNA binding to the subsequent anion exchange step **(Figure S3)**. The performance of two ion-exchangers - Q Sepharose and Fractogel DMAE, were next compared and both chromatography material showed similar plasmid DNA binding (**Figure S4)**. To confirm that the method was reproducible, three independent gravity flow purification columns at 0.2g cell weight scale were conducted using the optimized steps **(Figure 2A)**, and all three runs showed similar quality and quantity of recovered DNA **(Figure 2B)**. HPLC analysis confirmed no RNA was detectable and DNA was 85-88% supercoiled (SC) **(Figure 2C, Table 1)**.

**Table 1:** HPLC purity of 3 batches of plasmid DNA purified from replicate columns **(Figure 2)**. Percentage of each nucleic acid species was calculated based on the area under the curve.

| Samples | RNA<br>(AUC)<br>Peak 1 | Open<br>Circular<br>(AUC)<br>Peak 2 | Supercoiled<br>(AUC)<br>Peak 3 | %<br>Supercoiled |
| --- | --- | --- | --- | --- |
| Replicate 1 | 0 | 72,508 | 540,187 | 88% |
| Replicate 2 | 0 | 89,468 | 637,844 | 85% |
| Replicate 3 | 0 | 88,885 | 628,242 | 85% |

**Figure 2.**
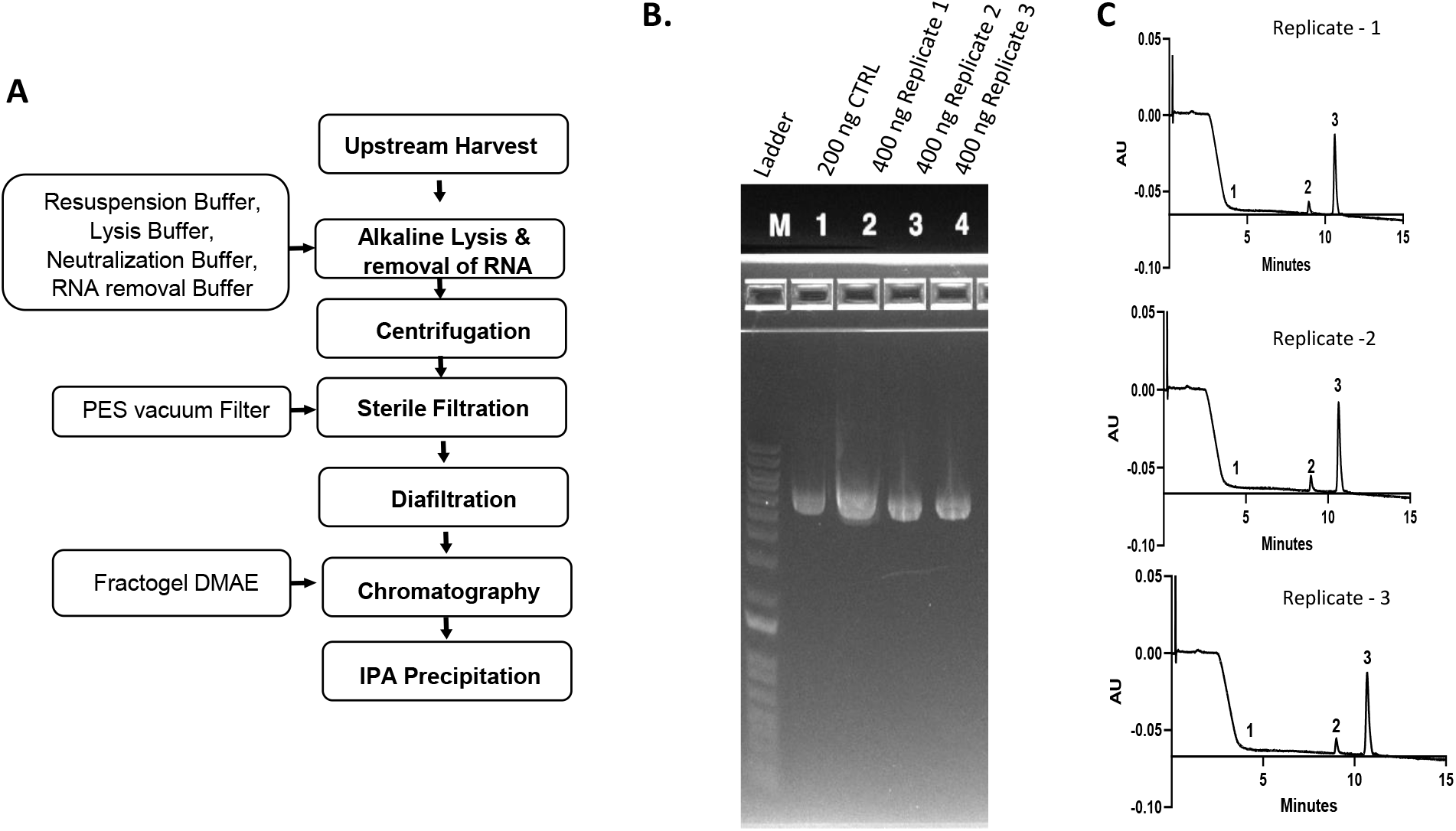
DNA purification optimization using gravity columns: **A)** Flow diagram of the production process. **B)** 5g-scale cell weight purification performed in replicate 0.5 mL DMAE Fractogel gravity columns and elution fractions analyzed by agarose gel electrophoresis: M, DNA Ladder; lane 1, control VEEV plasmid; lanes 2, 3, 4 show plasmid purified from replicate columns. **(C)** HPLC from three replicate columns. HPLC Peak-1, RNA; Peak-2, open circular DNA; Peak-3, supercoiled DNA.

### FPLC-based purification (5g-scale)

The gravity column work-flow was next adapted to an AKTA Pilot™ chromatography system. Three independent 5g-scale purification experiments were conducted **(Figure 3A)**. Following lysis, 125 mL lysate was clarified by centrifugation and sterile filtered. TFF allowed removal of small impurities and adjustment of conductivity to ∼44 mS/cm and pH ∼7.5 **(Table 2)**. The nucleic acid yield after TFF was ∼1 mg/gm cell weight. After TFF, ∼2 g paste equivalent was loaded onto Fractogel DMAE chromatography columns and 600 mM NaCl wash was used to remove residual RNA **(Figures 3A and 3B)**. Purified plasmid was eluted using 1 M NaCl and final isopropanol precipitation yielded ∼0.25 mg DNA/g cell paste. The final DNA product migrated as a single band on agarose gel **(Figure 3B)** and HPLC analysis confirmed >90% supercoiled pDNA and ≤2% RNA **(Figure 3C, Table 3)**.

**Table 2.** Representative pH, conductivity and yield parameters from the dilution and concentration TFF cycles performed at 5g-scale for plasmid purifications shown in **Figure 3**.

| Sample | Volume<br>(mL) | Conductivity<br>(mS/cm) | A260 | 260/280 | pH | Nucleic<br>Acid<br>(µg/mL) |
| --- | --- | --- | --- | --- | --- | --- |
| Lysate | 125 | 94 | 6.87 | 2.04 | 4.2 | 344 |
| TFF Cycle #1 | 125 | 83 | 3.34 | 2.08 | 4.5 | 167 |
| TFF Cycle #2 | 125 | 56 | 0.88 | 2.04 | 4.7 | 44 |
| TFF Cycle #3 | 125 | 47 | 0.4 | 2 | 5 | 20 |
| TFF Cycle #4 | 125 | 44 | 0.45 | 2 | 5.6 | 23 |
| TFF Cycle #5 | 125 | 44 | 0.8 | 2.01 | 7.5 | 40 |

**Table 3.** Purity of 3 batches of DNA purified using FLPC **(Figure 3B)** as determined by analytical HPLC. Percent supercoiled was calculated using area under the curve.

| Sample | RNA (AUC) Peak 1 | Open Circular (AUC) Peak 2 | Supercoiled (AUC) Peak 3 | % Supercoiled | % RNA |
| --- | --- | --- | --- | --- | --- |
| Replicate 1 | 75,822 | 324,382 | 2,808,219 | 90% | 2% |
| Replicate 2 | 0 | 132,359 | 2,492,440 | 95% | 0% |
| Replicate 3 | 0 | 100,702 | 2,149,392 | 96% | 0% |

**Figure 3.**
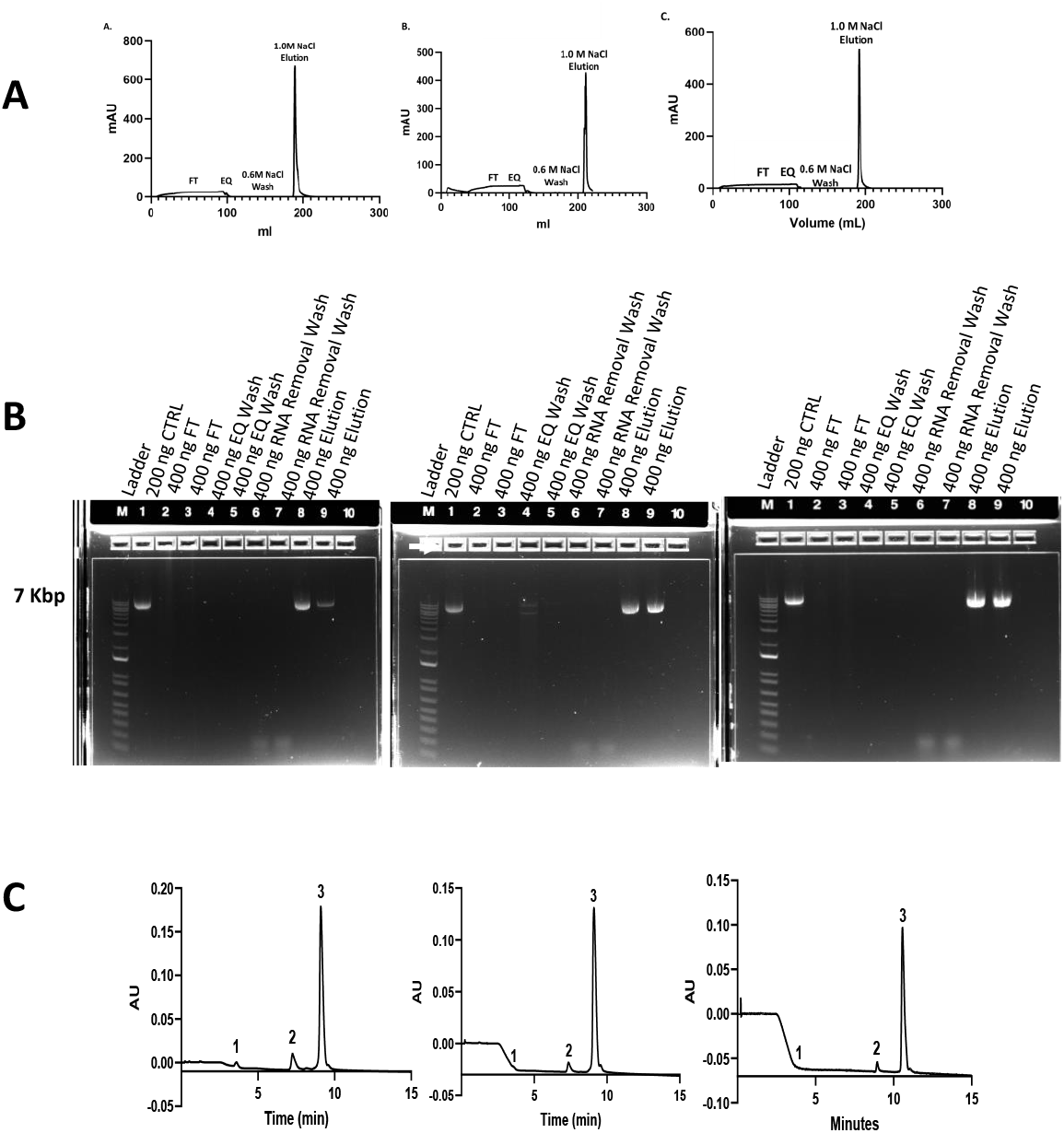
5g-scale FPLC-based purification: **(A)** Chromatograms of DMAE-Fractogel chromatography runs conducted in triplicate. **(B)** Plasmid analysis by agarose electrophoresis. Lanes: M, DNA Ladder; lane 1, Control VEEV plasmid; lanes 2, 3 flow-through fractions (FT); lanes 4, 5, equilibration wash fractions (EQ); lanes 6, 7, RNA wash fractions; lanes 8, 9 DNA in elution fractions. **(C)** HPLC analysis showing purity of three DNA batches. Peaks 1, 2, and 3 represent - RNA, open circular DNA and supercoiled DNA, respectively.

### Purification (50g-scale)

The purification protocol optimized at the 5g-scale, was scaled-up to 50g, equivalent to ∼1L fermentation culture **(Figure 4A)**. Alkaline lysis resulted in 1.2 L lysate that was cleared by a 3-step filtration process **(Figure S5)**, followed by 5 cycles of dilution and reconcentration using TFF to reach the final conductivity of 49 mS/cm and pH 7.7 (**Table 4)**. Post-TFF, the lysate contained ∼30 mg total nucleic acid which was purified on a 35 mL Fractogel DMAE column **(Figure 4B)**. The isopropanol precipitation step resulted in ∼7.5 mg DNA in 5 ml TE buffer (0.15 mg/g cell paste). The purified DNA was nearly 100% pure, containing >90% supercoiled isoform **(Figures 4C, Table 5)**. The final DNA product at ∼1 mg/ml tested negative for residual protein using NanoOrange Quantitation Kit (limit of quantitation ∼300 ng/ml).

**Table 4.** pH, conductivity and DNA yield from serial dilution and concentration TFF cycles, performed at 50g-scale.

| Sample | Volume (mL) | Conductivity (ms/cm) | A260 | A280 | 260/280 | pH | Nucleic Acid (µg/mL) |
| --- | --- | --- | --- | --- | --- | --- | --- |
| Lysate | 2700 | 87 | 3.01 | 1.36 | 2.22 | 5.1 | 113.5 |
| TFF Cycle #1 | 1200 | 84 | 2.99 | 1.35 | 2.21 | 5.1 | 149 |
| TFF Cycle #2 | 1200 | 71 | 1.57 | 0.72 | 2.18 | 5.6 | 79 |
| TFF Cycle #3 | 1200 | 60 | 1.01 | 0.45 | 2.23 | 7 | 50 |
| TFF Cycle #4 | 1200 | 55 | 0.61 | 0.27 | 2.25 | 7.5 | 31 |
| TFF Cycle #5 | 1200 | 49 | 0.48 | 0.21 | 2.2 | 7.7 | 24 |

**Table 5.**
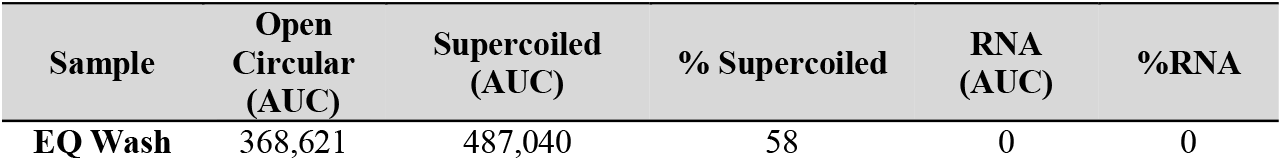
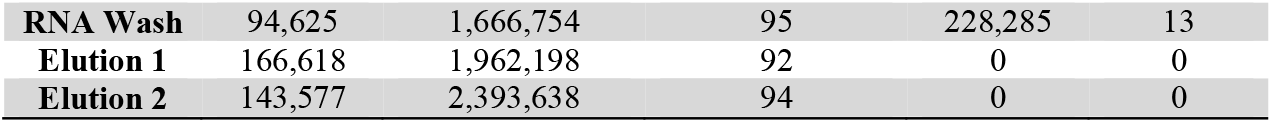
Purity of DNA fractions **(Figure 4C)** estimated using analytical HPLC. Percentage purity was calculated based on the area under the curve.

**Figure 4.**
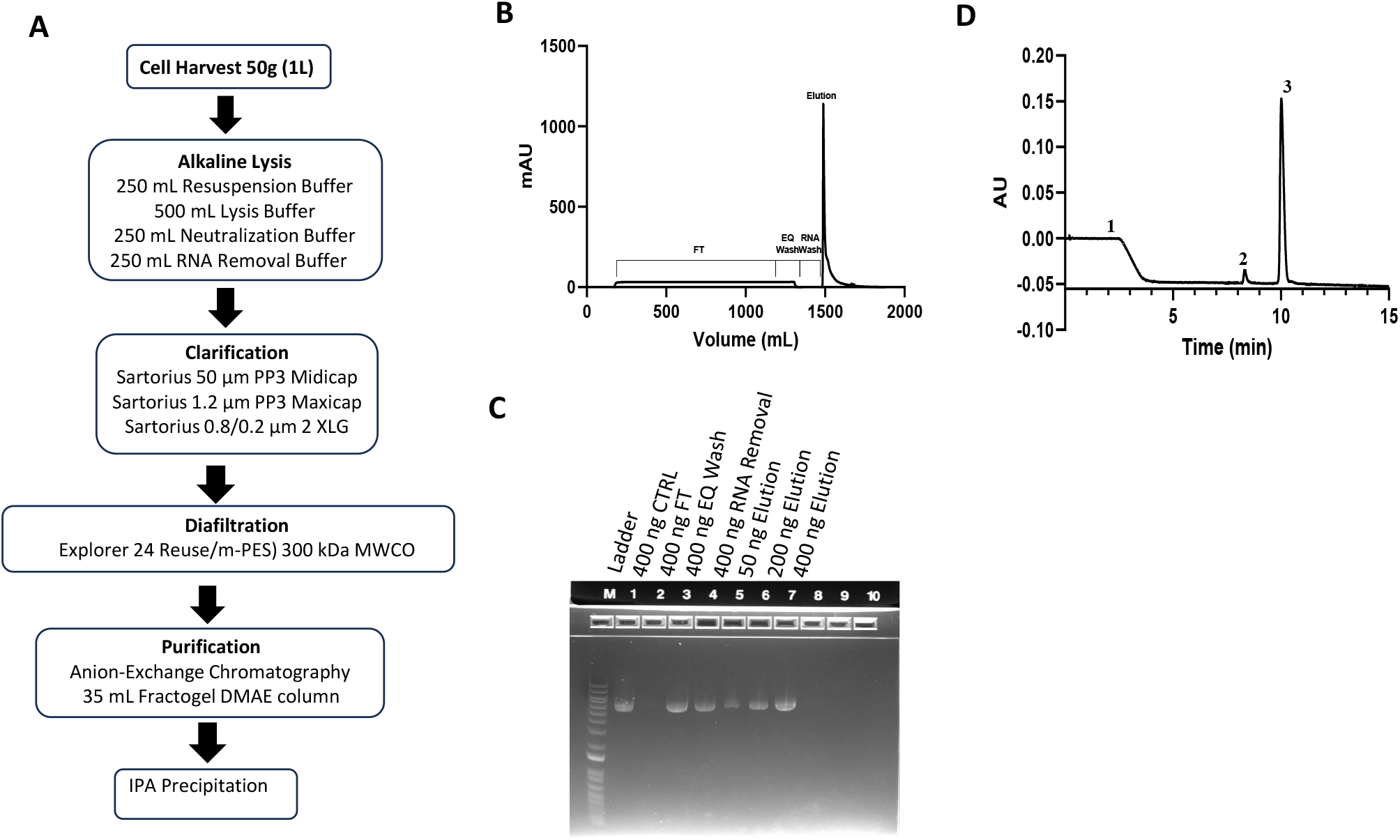
Plasmid DNA production at 50g-scale: **(A)** Workflow for down-stream purification at 50g-scale. **(B)** Elution chromatogram of 35 mL DMAE Fractogel column. **(C)** Agarose gel analysis of chromatography fractions. Lanes: M, DNA Ladder; lane 1, control VEEV plasmid; lane 2, flow-through (FT); lane 3, equilibration buffer wash (EQ); lane 4, RNA wash; lane 5, 6, 7 elution fractions. **(D)** HPLC analysis of the elution fraction. Peaks labelled 1, 2, 3 represent - RNA, open circular DNA, and supercoiled DNA, respectively

## Discussion

COVID-19 pandemic was a pivot point, dispelling the fallacy that low- and middle-income countries (LMIC) can rely on western infrastructure for timely production and delivery of life-saving vaccines. Pure plasmid is the starting point for the development of DNA vaccines, RNA vaccines, non-viral gene therapy trials, early-stage transient transfection-based trial material, cell-free translation products and genetic manipulation of micro-organisms. Early-phase clinical trials require 1–5 g purified plasmid DNA, manufactured under cGMP conditions, and pandemic response for human or veterinary use, would require kilogram quantities of high-quality plasmid. Contract manufacturing and in-licensing of commercial technologies that meet international regulatory specifications can be expensive [29]. Open access nucleic acid purification protocols such as the one described here could enable timely access to locally sourced DNA vaccines.

The DNA purification schema described here needs to be further optimized for cGMP manufacture. Fermentation using alternative feed strategies or switching to a defined synthetic medium can increase the biomass density during fermentation where OD_600_ >100 can be achieved. Growth medium conditions can also be modified to increase the percent supercoil yield [34, 35]. Bacterial harvest was performed here using centrifugation which is not ideal for high volume fermentation. An ultra-filtration based method using 1.2 µm pore size hollow fiber membrane can be used to exchange the culture medium directly into Resuspension Buffer, allowing cell lysis to proceed immediately following fermentation. Manufacturing cost can be further economized by incorporation of vacuum or CO_2_ based separation of the flocculant material which would reduce bioburden before sterile filtration [36].

High percentage of supercoiled DNA is required for efficient transfection and elicitation of a productive immune response in mammals [37, 38]. *E. coli* typically produces negatively charged supercoiled DNA which requires elastic energy to maintain its shape. Positively charged ions can screen the negative charges, affecting supercoiling, and higher salt concentrations keep the DNA compacted allowing the elastic energy to dominate over electrostatic energy [39, 40]. We maintained high salt concentration ∼400 mM during the high-shear operations, particularly TFF, resulting in a higher percentage of supercoiled plasmid DNA. Another key factor that affected supercoiling was the time-period between lysis and isopropanol precipitation. Halting purification mid-process was detrimental and shortening the process time led to improved supercoiled plasmid yield.

Final purification was performed using DMAE Fractogel anion exchanger where a 600 mM NaCl wash removed residual RNA contaminants and plasmid DNA was eluted with 1 M NaCl. Concentration and transfer of the plasmid to TE buffer was performed using isopropanol precipitation which could also be replaced by TFF. Milli-Q grade water was used here however utilizing ultra-pure “Water For Injection (WFI) and introducing Triton X-100 during chromatography can reduce the residual endotoxin [41]. At 50 g (1 L) scale, our process yielded ∼7.5 mg pDNA or 0.15 mg/g paste which was ∼40% less than the target yield (0.25 mg/g paste). One reason for this difference could be the use of Fractogel DMAE that has a pore size ∼800 Å and is smaller than the size of supercoiled plasmid DNA. Increasing resin bed volume could allow more efficient capture of plasmid during chromatography. Alternatively, CaptoCore™ based resins can be used to efficiently fractionate supercoiled DNA [42]. Notwithstanding the need for further improvement, we demonstrated the feasibility of producing ∼99% pure plasmid DNA with >90% supercoiled content which is the pre-requisite for use in a DNA vaccine [28].

## Acknowledgements

We thank Dr. Jay W. Hooper, Virology Division, United States Army Medical Research Institute of Infectious Diseases, Fort Detrick for manuscript review and guidance and plasmid starting material. Funding for this work was provided by Defense Threat Reduction Agency (DTRA) Joint Science and Technology Office (JSTO).

## Authors contributions

TLWilson: Methodology, analysis, writing and editing; SD: Concept generation, writing, editing, supervision; MH: Methodology, investigation, analysis; CP: Methodology; FK: Methodology, SZ: Funding and Project Management.

## Disclaimer

The opinions or assertions contained herein are the private views of the authors and are not to be construed as official or as reflecting the true view of the Department of the Army, the Department of Defense, or the US Government. The work has been cleared for publication by the Walter Reed Army Institute of Research and the Defense Threat Reduction Agency Joint Science and Technology Office.

## Funding

Funding for this work was provided by Defense Threat Reduction Agency (DTRA) Joint Science and Technology Office (JSTO).

## Notes

### Competing Interest Statement

The authors have declared no competing interest.

