## Supplementary Figures for "RNAse-free manufacture of Venezuelan Equine Encephalitis Virus (VEEV) plasmid DNA vaccine"

**Figure S1:** Sequencing results of fermentation product confirming plasmid identity.

**Processed Sample Alignment with VEEV Sequence (BGH Primer)**

```
Sample      ------------------TTATATCCGGTACCATGATTGCCNCTGCGACAGATCAGTATA	42
VEEV        CGTGCACGTGCCCAGCGGCACAGCCACACTGAAGGAAGCCGCCGTGGAGCTGACCGAGCA	5580
                                    *       * **   *  *   **     * *   *

Sample      GAGCCGCGCCACCATCCACTTCAGCACCGCCAACATCCACCCCGAGTTCAGGCTGCAGAT	102
VEEV        GGGCAGCGCCACCATCCACTTCAGCACCGCCAACATCCACCCCGAGTTCAGGCTGCAGAT	5640
            * ** *******************************************************

Sample      TTGCACCAGCTACGTGACATGCAAGGGCGACTGCCACCCCCCTAAGGACCACATCGTGAC	162
VEEV        TTGCACCAGCTACGTGACATGCAAGGGCGACTGCCACCCCCCTAAGGACCACATCGTGAC	5700
            ************************************************************

Sample      CCACCCCCAGTACCACGCCCAGACCTTCACAGCCGCCGTGTCCAAGACAGNNTGGACCTG	222
VEEV        CCACCCCCAGTACCACGCCCAGACCTTCACAGCCGCCGTGTCCAAGACAGCCTGGACCTG	5760
            **************************************************  ********

Sample      GCTGACCAGCCTGCTGGGCGGCAGCGCCGTNNNCTTCATCATCGGCCTGGTGCTGGCCAC	282
VEEV        GCTGACCAGCCTGCTGGGCGGCAGCGCCGTGATCATCATCATCGGCCTGGTGCTGGCCAC	5820
            ******************************   * *************************

Sample      CATCGTGGCCATGTACGTGCNGACCAACCAGAAACACANNNNANNNNNNNNNNNN-----	337
VEEV        CATCGTGGCCATGTACGTGCTGACCAACCAGAAACACAACTGATGAAGATCTACGTATGA	5880
            ******************** *****************    *
```

**Figure S2.** **Effect of CaCl_2_ addition on RNA content:** 10g cell paste was subjected to alkaline lysis and half was treated with 1M Calcium Chloride. The material was clarified by centrifugation (8000xg) for 10 min, filtered on a 0.45 µm membrane, followed by 0.2 µm sterile filter (Millipak Gamma). Diafiltration was performed using Explorer-12 Hollow Fiber Membrane (300 kDa MWCO). After five cycles of buffer exchange in the TFF, retentate aliquots were collected and evaluated by agarose gel. The results showed significant reduction in RNA, following dialfiltration and CaCl_2_ addition. Lanes: M, Ladder; Lane 1, VEEV plasmid positive control; lane 2, TFF retentate post Cycle-1; lane 3, Cycle 2; lane 4, Cycle 3; lane 5, Cycle 4; lane 6, Cycle 5; lane 7, permeate post Cycle-3; lane 8, permeate post Cycle-5.

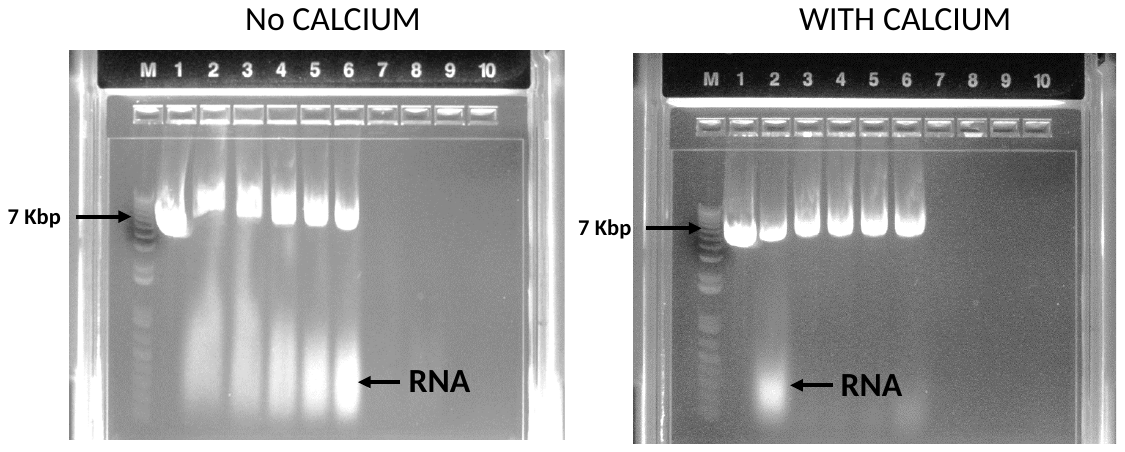

**Figure S3.** **Effect of diafiltration on the recovery of pDNA:** At 5g-scale, cells were subjected to alkaline lysis, cleared by centrifugation (8000xG) and sterile filtrated using 0.22 µm filter (Nalgene). The clarified lysate was subjected to subsequent rounds of dilution and concentration by TFF. Aliquots of the retentate from each cycle (5 ml) were tested for binding to 0.5 mL Fractogel DMAE gravity columns. The columns were washed with 2 ml 0.6 M NaCl, DNA eluted in 2 ml 1M NaCl, precipitated with IPA (1:1 v/v) and recovered by centrifugation (10,000xG) for 30 min at 4°C. DNA pellets were resuspended in 100 µL TE Buffer and analyzed by agarose gel. The gel shows optimal binding to Fractogel DMAE after 4-5 cycles of diafiltration. **(A)** Table shows the pH and conductivity changes during TFF. **(B)** Agarose gel analysis. Lanes M, DNA Ladder; lane 1, control plasmid (200 ng); lane 2, DNA eluted from the DMAE after TFF Cycle 1; lane 3, TFF Cycle 2; lane 4, TFF Cycle 3; lane 5, TFF Cycle 4; lane 6, DNA eluted from DMAE after TFF Cycle 5.

**A	B**

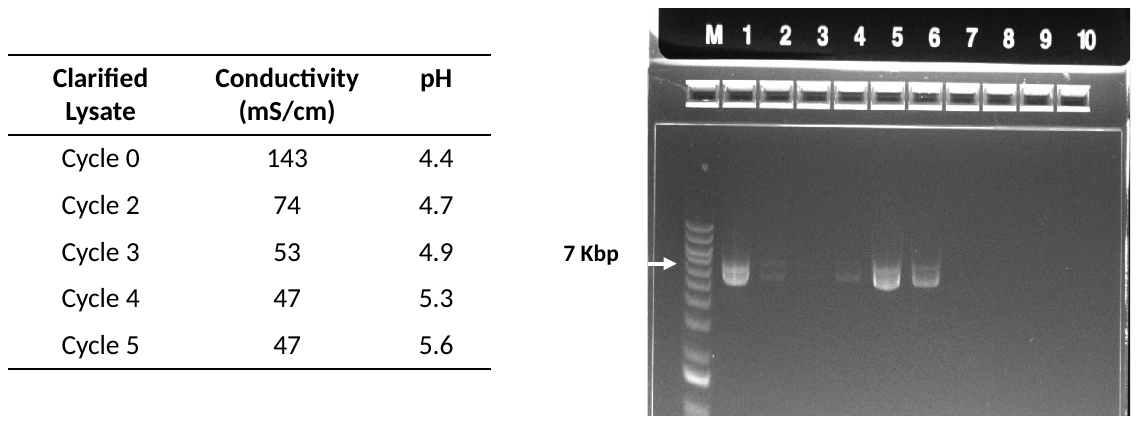

**Figure S4.** **Comparison of Q Sepharose and DMAE Fractogel:** At 5 g scale, cell paste was subjected to alkaline lysis, cleared by centrifugation (8000xG) and sterile filtered using 0.22 µm filter (Nalgene). The clarified lysate was diluted and concentrated five times using TFF (MidiKross Hollow Fiber Membrane; 300 kDa MWCO). Aliquots (5 mL) were collected and loaded onto 0.05 mL, 0.25 mL and 0.50 mL of Q Sepharose or Fractogel DMAE. Columns were washed with 0.6 M NaCl (2 ml) and eluted with 1 M NaCl (2 ml). Samples were precipitated by adding IPA (1:1 v/v), and DNA was harvested by centrifugation (10,000x rpm for 30 min at 4°C). The pellets were air-dried for 30 min and suspended in 100 µL TE. The Fractogel DMAE resin showed higher yields. **(A, B)** Lanes: M, DNA Ladder; lane 1, control plasmid (200 ng); lanes, 2, 3, 4, flow-through fractions from 0.05, 0.25 and 0.5 mL resin; lanes 5, 6, 7, RNA wash from 0.0, 0.25, 0.5 mL resin; lanes 8, 9, 10, elution fractions from 0.05, 0.25, 0.5 mL resin, respectively. **(C)** Table shows DNA yield and recovery from each column.

**A**	**B**	**C**

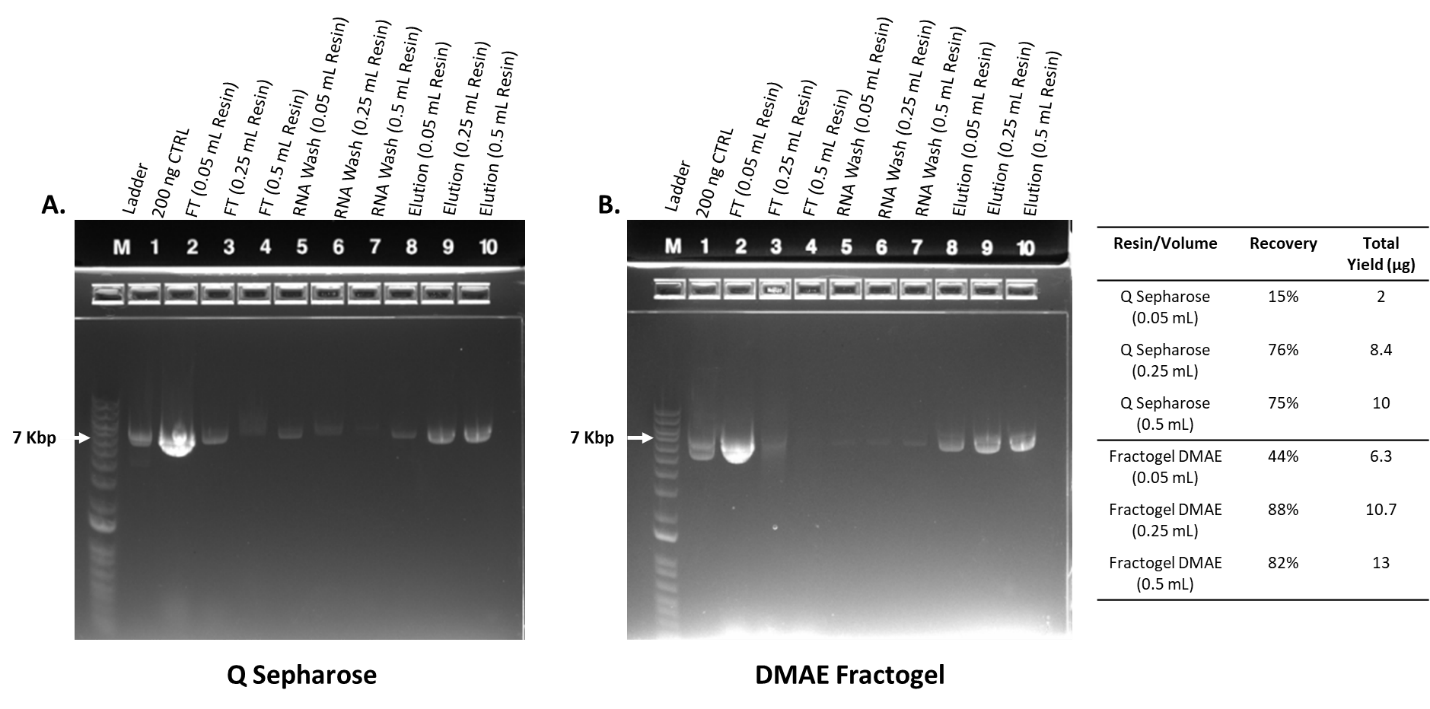

**Figure S5.** **Clarification of lysate by serial filtration at 50 g scale.** Filtration was performed on the lysate using Sartopure-PP3 Midicap (50-micron) **(A);** followed by Sartopure-PP3 Maxicap (1.2 micron) **(B);** and finally Sartopore-2XLG (0.8/0.2-micron) **(C)**. Each filter was equilibrated and washed with the diafiltration buffer before and after filtering the lysate using a peristaltic pump (60 mL/min). A final volume of 2.7 L was clear and used for diafiltration.

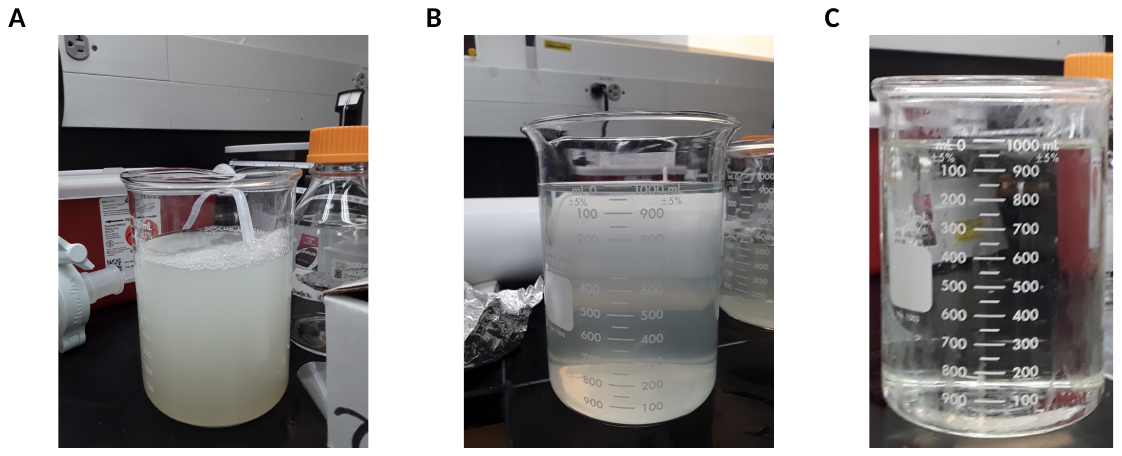
